# Patient-derived glioblastoma neurosphere cultures differentially express nicotinic acetylcholine receptors depending on ambient choline

**DOI:** 10.1101/2023.04.06.535046

**Authors:** Elena A. Gondarenko, Diana V. Mazur, Marina Masliakova, Yana A. Ryabukha, Igor E. Kasheverov, Victor I. Tsetlin, Denis S. Kudryavtsev, Nadine V. Antipova

## Abstract

Glioblastoma multiforme (GBM) is the most aggressive type of brain cancer with a poor prognosis. GBM cells, developing in the environment of neural tissue, often exploit neurotransmitters and their receptors to promote their growth and invasion. Nicotinic acetylcholine receptors (nAChRs) play a crucial role in the central nervous system signal transmission, are widely represented in the brain, the GBM cells expressing several subtypes of nAChRs which are suggested to transmit signals from neurons, thus promoting tumor invasion and growth. Functional α1*, α7 and α9 nAChRs are demonstrated on several patient-derived GBM neurosphere cultures and U87MG cell line using neurotoxins and fluorescent calcium assay. Selective α1*, α7 and α9 nAChR antagonists stimulated cell growth in presence of nicotinic agonists. Choline, normally present in blood, is capable of activating α1*, α7 and α9 nAChR subtypes, mediates the antagonist’s influence on cell proliferation. Several cultivating conditions have been shown to directly change sensitivity of primary GBM lines to nAChR ligands. Thus, results of *in vitro* testing of nAChR ligands on GBM lines should be interpreted and reviewed in cell culture conditions-aware manner.

## INTRODUCTION

Glioblastoma multiforme (GBM) is the most aggressive and lethal primary tumor of the central nervous system (CNS) with the patient median survival of 15 months from the time of diagnosis [1]. Unfortunately, current standard-of-care, including surgical resection, radiotherapy and oral administration of DNA-alkylating drug temozolomide, show limited efficacy in GBM treatment.

The Cancer Genome Atlas (TCGA) proposed four main GBM subtypes based on their genomic features: proneural, mesenchymal, neural, and classical [2]. Several studies came to results which can be used to characterize various subtypes of the tumor [3–5]. Recent data show that a proneural-mesenchymal transition (PMT) may occur at different stages of the disease. Tumor transformation into a mesenchymal subtype GBM stem cells is associated with more aggressive phenotypes, treatment resistance and tumor recurrence [6,7].

The microenvironment of GBM cells also plays an important role in the development, recurrence and resistance of brain cancer. GBM cells undergo dynamic and reversible transitions in response to changes in the microenvironment, which in turn changes the spatial and temporal phenotypic heterogeneity of the tumor [8]. Numerous studies show that interaction with neurons, as well as direct and indirect consequences of neuronal activity, can influence tumor progression [9,10].

It has been shown that neurons affect tumor growth and progression by forming functional synapses with glioma cells [11,12]. For instance, GABA-mediated synaptic transmission inhibits GBM growth, whereas active release of glutamate increases cell proliferation [13]. Other signaling molecules transmitted between glioma cells and normal neurons are brain neurotrophic factor, neuroligin-3 and dopamine [14]. The cholinergic projections are also known to innervate regions where GBM develops, releasing neurotransmitter acetylcholine (ACh) [15].

Several subtypes (mainly α1, α7, α9 and α10)) of nicotinic acetylcholine receptors (nAChRs) effectively permeate calcium ions [16]. It has been confirmed that GBM cells express certain types of nAChRs and the functionality of these receptors has been demonstrated [15,17]. The expression of the majority of nAChR genes in GBM cells is at a lower level compared to normal. In primary cell lines most frequently expressed nAChR subunits mRNAs are CHRNA5 (which encodes α5 subunit) and CHRNA9 (α9), and at a lower level – CHRNA7 (α7) and CHRNA10 (α10). In the model lines U87MG the expression of muscle nAChR subunits CHRNA1 (α1) and CHRNB1 (β1) was also shown [15,17].

Intracellular calcium has been implicated in various signaling pathways that affect tumor progression. Several studies suggest that GBM proliferation, migration and invasion are changed in the presence of nAChR ligands. These effects are predominantly mediated by α7 and α9 nAChRs. Some GBM model lines such as U251 proliferate in the presence of AChR agonist acetylcholine [15]. Complementary to that, 10^-7^ M nicotine increases the proliferation of the U87MG and GBM lines. Nicotine effect is blocked by both α-bungarotoxin (α-Bgt) and mecamylamine (MLA), which un-selectively inhibit muscle nAChR, α7 and α9 nAChRs, as well as by α7-selective antagonist α-conotoxin derivative ArIB [V11 L; V16D] and α9-selective antagonist α-conotoxin RgIA [17].

Furthermore, GBM can easily develop radio- and chemoresistance_due to the extreme heterogeneity of these tumors [18]. Different GBM cell subpopulations retain distinct genetic profiles, biomolecular markers and functional properties. In this regard, it is crucial to choose the right models to study molecular and biochemical features and test the effects of potential therapeutic substances. Patient-derived cancer stem cell enriched cultures (CSCs) that are grown in serum-free media are the most accepted standard for studying the biology of GBM in vitro [19]. However, the same primary cultures grown under standard *in vitro* conditions with serum-supplemented media have gene expression profiles that are dramatically different in either SSCs or the original tumors [20].

In this paper we explored cholinergic signaling carried out by nicotinic receptors in primary GBM lines and model U87MG GBM line using a range of methods: from real time PCR and fluorescent calcium imaging to AlamarBlue proliferation assay. We studied the effect of cultivation conditions on primary cultures for the expression and functional activity of distinct nAChRs, as well as the effect of nAChR ligands on cell viability in various growing options (Fig. 1). Epibatidine activates all nAChR subtypes except α9, for which it is known to be an inhibitor. In order to determine which specific nAChR subtypes activation is associated with changes in GBM cell proliferation, the α1 and α7 subunit antagonists azemiopsin and α-conotoxin [A10L]PnIA were added to cells along with epibatidine or without any externally added agonists.

**Figure 1.**
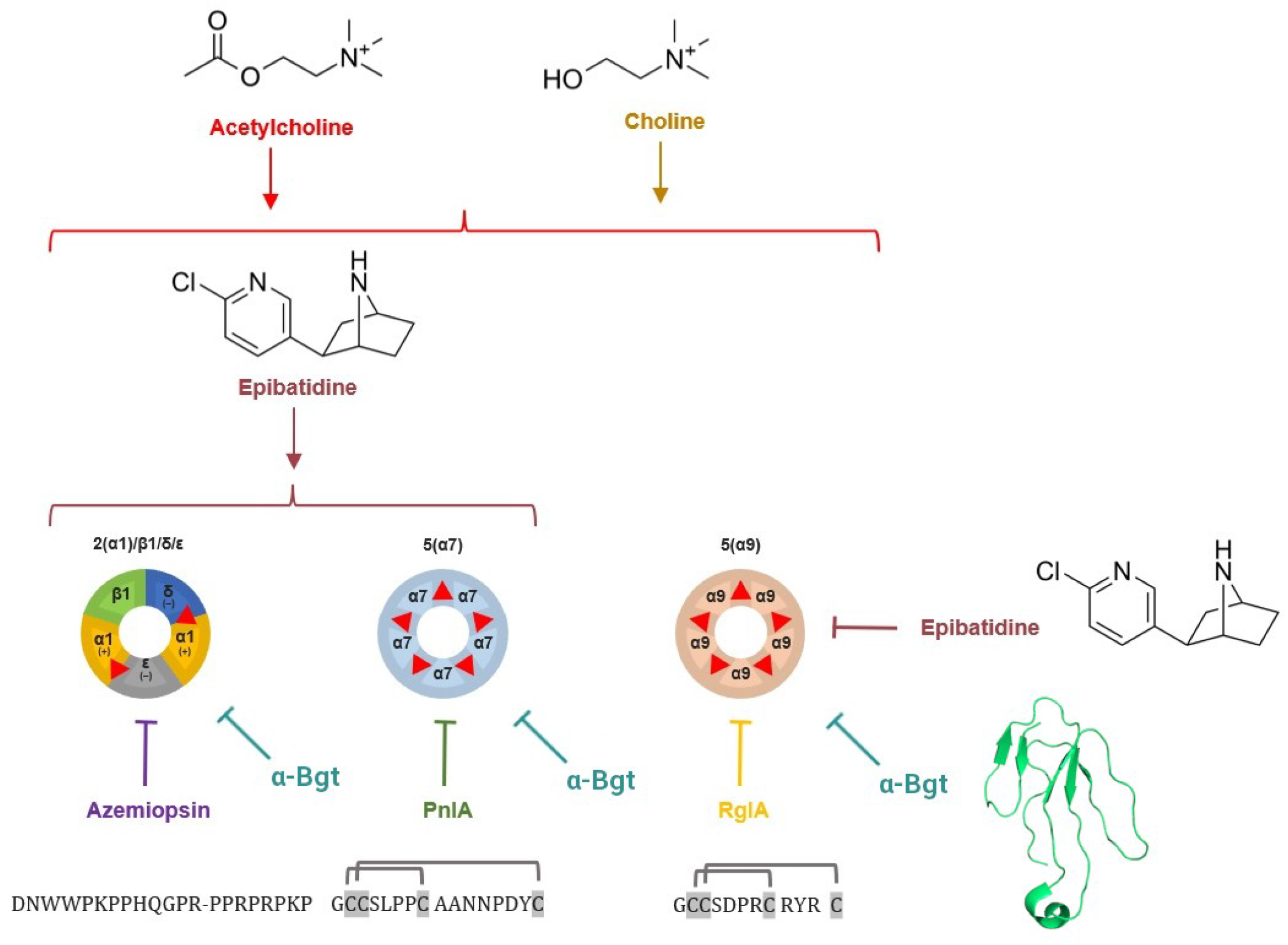
Subtypes of nAChR covered in this article – heteromeric muscle α1β1δε receptor and neuronal α7 and α9 homopentameric receptors. Orthosteric binding sites for agonists and competitive antagonists are illustrated as red triangles. Orthosteric agonists: choline, acetylcholine and epibatidine shown as chemical formulae, peptide neurotoxins azemiopsin, [A10L]PnIA and RgIA shown by their respective sequences, small protein α-bungarotoxin is shown by ribbon representation of secondary structure.

## MATERIALS AND METHODS

### Ethics statement

Tumor tissue samples from GBM patients were taken during surgery. The diagnosis was established on the basis of histological analysis according to the classification of brain tumors proposed by the World Health Organization (WHO). All procedures performed in studies involving human participants were in accordance with the ethical standards of the institutional and/or national research committee and with the 1964 Helsinki Declaration and its later amendments or comparable ethical standards. The study was approved by the ethical committee of the N.N. Burdenko National Medical Research Center of Neurosurgery, protocol #176 from August 29, 2019. All the patients gave written informed consent for their participation. Tissues were processed in the research laboratory after de-identification of the samples.

#### 2.1 Media for the cultivation of GBM cells

Medium I: DMEM/F12 (Sigma–Aldrich), containing 2 mM glutamine, 10% fetal bovine serum (FBS) (Gibco, Thermo Fisher Scientific), 1% Penicillin-Streptomycin solution (Sigma–Aldrich).

Medium II: DMEM/F12 (Sigma–Aldrich), containing 2 mM glutamine, 2% NS-21 (MACS NeuroBrew-21 supplement (Miltenyi Biotec)), 20 ng ml^-1^ basic fibroblast growth factor (bFGF; Sigma– Aldrich), 20 ng ml^-1^ epidermal growth factor (EGF; Sigma–Aldrich), 1% Penicillin-Streptomycin solution (Sigma– Aldrich).

Medium III: DMEM/F12 (Sigma–Aldrich), containing 2 mM glutamine, 2% NS-21 (MACS NeuroBrew-21 supplement (Miltenyi Biotec)), 20 ng ml^-1^ basic fibroblast growth factor (bFGF; Sigma– Aldrich), 1% Penicillin-Streptomycin solution (Sigma–Aldrich).

Medium IV: DMEM/F12 (Sigma–Aldrich), containing 2 mM glutamine, 2% NS-21 (MACS NeuroBrew-21 supplement (Miltenyi Biotec)), 1% Penicillin-Streptomycin solution (Sigma–Aldrich).

Medium V: DMEM/F12 (Sigma–Aldrich), containing 2 mM glutamine, 2%, 1% Penicillin-Streptomycin solution (Sigma–Aldrich).

#### 2.2. Cell culture

All cells were cultured at 37 °C in a humidified atmosphere with 5% CO2. bFGF and EGF were added twice weekly, and the culture medium was changed every 5-10 days. To attach the cells to the glass or plastic, the surface was preincubated overnight at room temperature with a solution of laminin in PBS (1:200). Primary culture spheres were dissociated using StemPro Accutase (Thermo Fisher Scientific). Cells of the U87MG cells were dissociated using Trypsin-Versen solution (Paneco).

#### 2.3. RNA Isolation and qPCR

RNA was isolated using the ExtractRNA kit (Evrogen). RNA concentration was determined using a Nanodrop One C spectrophotometer (Thermo Fisher Scientific). cDNA was synthesized using the MMLV reverse transcription kit (Evrogen) according to the manufacturer’s protocol. qPCR was performed on LightCycler 96 (Roche) with qPCRmix-HS SYBR reagent (Evrogen). The cycling conditions were 95 °C for 150 s, followed by 45 cycles of 95 °C for 20 s, 60 °C for 20 s, and 72 °C for 20 s. Data were collected using LightCycler Software (version 4.1). 18S RNA was used as an intermediate control. The primer sequences are provided in Supplemental Table 1.

#### 2.4. Calcium Imaging

Primary cultures of GBM cells, as well as a U87MG cells were grown in 96-well plates in media I and II at 37 ° C, 5% CO2 atmosphere, 100% humidity. Before calcium imaging began, the growing medium was replaced with extracellular buffer (140 mM NaCl, 2 mM CaCl2, 2.8 mM KCl, 4 mM MgCl2, 20 mM HEPES, 10 mM glucose, pH 7.4) and then each well was loaded with a cell-permeant 5 μM Fluo-4AM solution (ThermoFisher Scientific) for 40 minutes. The fluorescent dye solution was further removed and cells were kept in an extracellular buffer for 1 hour. Ca^2+^ dynamics were recorded in the presence of different nAChR ligands using an Olympus IX71 epifluorescent microscope to observe fluorescence at 535 ± 10 nm. Data analysis was performed using open-source ImageJ Fiji software where changes in fluorescent intensity per cell before and after the nAChR modulators exposure were calculated. The response of at least 5 cells was measured.

#### 2.5. Patch-clamp

Cells were grown on laminin-covered glass placed to 24-well plates under the same conditions described in “Calcium imaging” subsection. To conduct the electrophysiological recording of the α7 nAChR-mediated macroscopic currents whole-cell patch clamp was set up as follows. Cells attached to the glass were transferred to the bath filled with extracellular electrophysiology solution (140 mM NaCl, 2 mM CaCl2, 2.8 mM KCl, 4 mM MgCl2, 20 mM HEPES, 10 mM glucose, pH 7.4). Microelectrode pipette was filled with intracellular buffer solution (140 mm CsCl, 6 mm CaCl2, 2 mm MgCl2, 2 mm MgATP, 0.4 mm NaGTP, 10 mm HEPES/CsOH, 20 mm BAPTA/KOH). Microelectrode was attached to cell membrane until resistance of at least 1 GOhm was reached. After establishing the gigaseal cell membrane was ruptured using suction and the recordings were made. A typical experiment consisted of 1 s cell wash with control extracellular buffer, then bath solution was changed to extracellular solution with 1, 10 or 100 μM of α7 nAChR-selective agonist PNU-282987 supplemented with 10 μM of α7 nAChR-selective positive allosteric modulator PNU-120596 for 5 s followed by 5 s wash-out with control extracellular buffer.

#### 2.6. Cell proliferation assay

Cells were plated in a 96-well plate at a density of 6,000 cells per well in 150 μl medium. The number of cells was assessed using AlamarBlue reagent (Thermo Fisher Scientific). The fluorescence was measured using a Fusion α-FP HT Universal Microplate Analyzer (PerkinElmer) with an excitation filter for 535 nm and an emission filter for 620 nm. The measurements were taken on day 5.

## RESULTS

### Transcriptomics database analysis

The Ivy Glioblastoma Atlas Project (Ivy GAP) database was used for the analysis. A matrix with RNA sequencing data of the nAChR genes CHRNA1-7, CHRNA9-10, CHRNB1-4, CHRND, CHRNE, CHRNG and mAChR genes CHRM1-5 was obtained from the database (Fig. 2).

**Figure 2.**
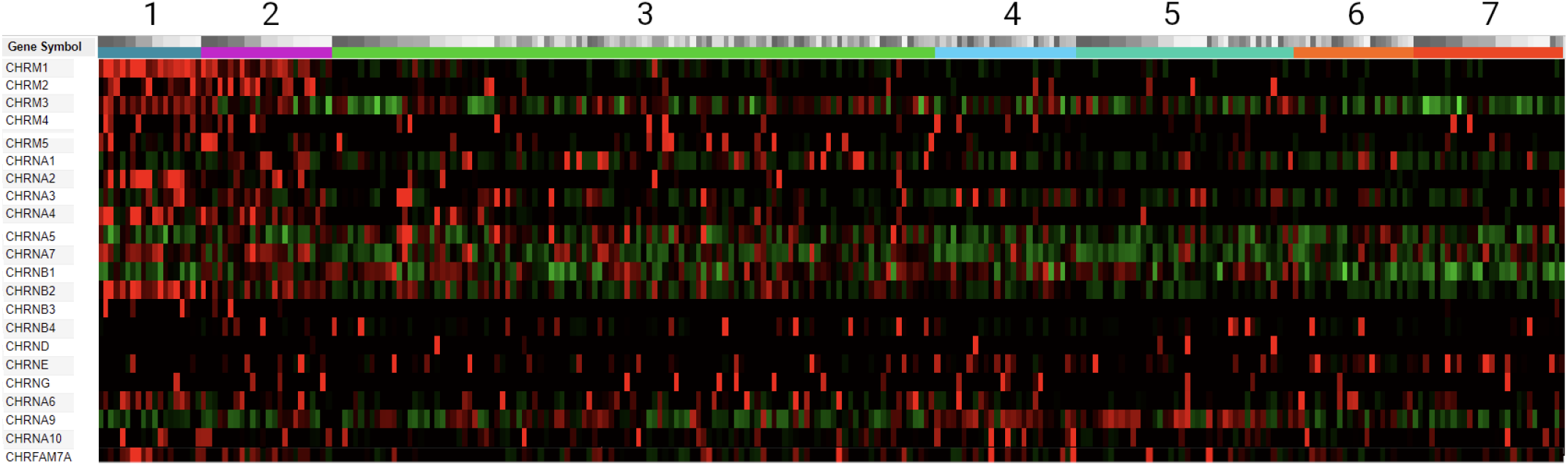
Differential expression of acetylcholine receptor genes (RNA-Seq mRNA data from Ivy GAP). Gene expression values are divided by regions and varieties of GBM from which samples were taken: 1) leading edge samples (blue-green); 2) infiltrating tumor samples (deep magenta); 3) cellular tumor samples (electric green);4) perinecrotic zone samples (picton blue);5) pseudopalisading cells around necrosis zone samples (caribbean green); 6) hyperplastic blood vessels in cellular tumor samples (orange); 7) microvascular proliferation samples (electric orange). The expression level is compared relative to the z-score value. A z-score value other than zero indicates the standard deviation from the average expression value of this gene. Therefore, a positive and negative z-score indicates increased and decreased gene expression, respectively. On the given heatmap the high z-score value is indicated in red, the z-score value below zero depicted by green, and the z-score value equal to or close to zero is represented by black.

Gene expression data were divided to seven different zones. The zones reflect the relationship between the anatomical features of GBM and the gene expression profile. As can be seen in Fig. 2, significant differences in the expression level of acetylcholine receptor genes are observed between samples for several genes. In particular, the CHRM1, CHRNA2, CHRNA4, CHRNA6, CHRNA7, CHRNB2 genes are increased in samples from the tissue of the anterior edge of the tumor. The expression of the CHRNA9 gene is reduced in the same region but increased in perinecrotic zone and pseudopalisading cells (Fig. 2). It is worth to note that progressing glioblastoma lesions are characterized by markedly elevated choline levels [21,22]. Thus, expression profile of choline-sensitive muscle (α1β1δε), α7 and α9α10 nAChRs are of a particular interest [23–25].

### Real-time qPCR detection of nAChR subunits

The results of RNA-Seq database analysis from Ivy GAP demonstrate differential expression of choline-sensitive nAChRs (α7 and α9) in different morphological zones of the tumor. Apart from brain lesions, choline is present in various media, e.g blood plasma concentration of choline in recent study was estimated using LC-MS/MS as high as 11.4 ng/μL, which corresponds to ~ 100 μM [26], although NMR data allow to estimate choline concentrations of up to 28.9 μM of choline in blood serum of healthy individuals [27]. It was decided to study nAChR genes expression by the patient-derived GBM neurosphere cultures using real-time PCR after culturing cells in two types of medium: without adding fetal bovine serum (but with adding bFGF and EGF growth factors and NS-21 neuronal vitality supplementation) and with adding FBS. Most of the lines studied showed higher levels of nAChR gene expression after cultivation under serum-free conditions compared with cultivation under serum conditions (Fig. 3 A). Interestingly, α5, α7 and α9 nAChR subunits cDNAs were detected in all of the lines studied, both under serum and serum-free conditions (Fig. 3 B and C). The U87MG cell line showed similar expression profiles in both media variants. The 011 pro-neuronal line showed almost all nAChR subunits genes expression at the mRNA level (except for α4, and β2 in serum-free medium), whereas the 019 pro-neuronal line expressed α1, α5, and α9 nAChR genes in serum-free conditions, while α7 nAChR gene expression was activated in medium with FBS in addition to other nAChR subunits. Interestingly, the response of pro-neuronal lines to cultivation in medium with FBS manifested itself in the emergence of new subunits expression that were not observed in serum-free conditions. Cell lines of the mesenchymal subtype showed the presence of α5, α7, and α9 nAChR subunits in both media variants. Mesenchymal subtype line 022 demonstrated additional expression of α1, α3, α6, α10, β2 nAChR subunits genes.

**Figure 3.**
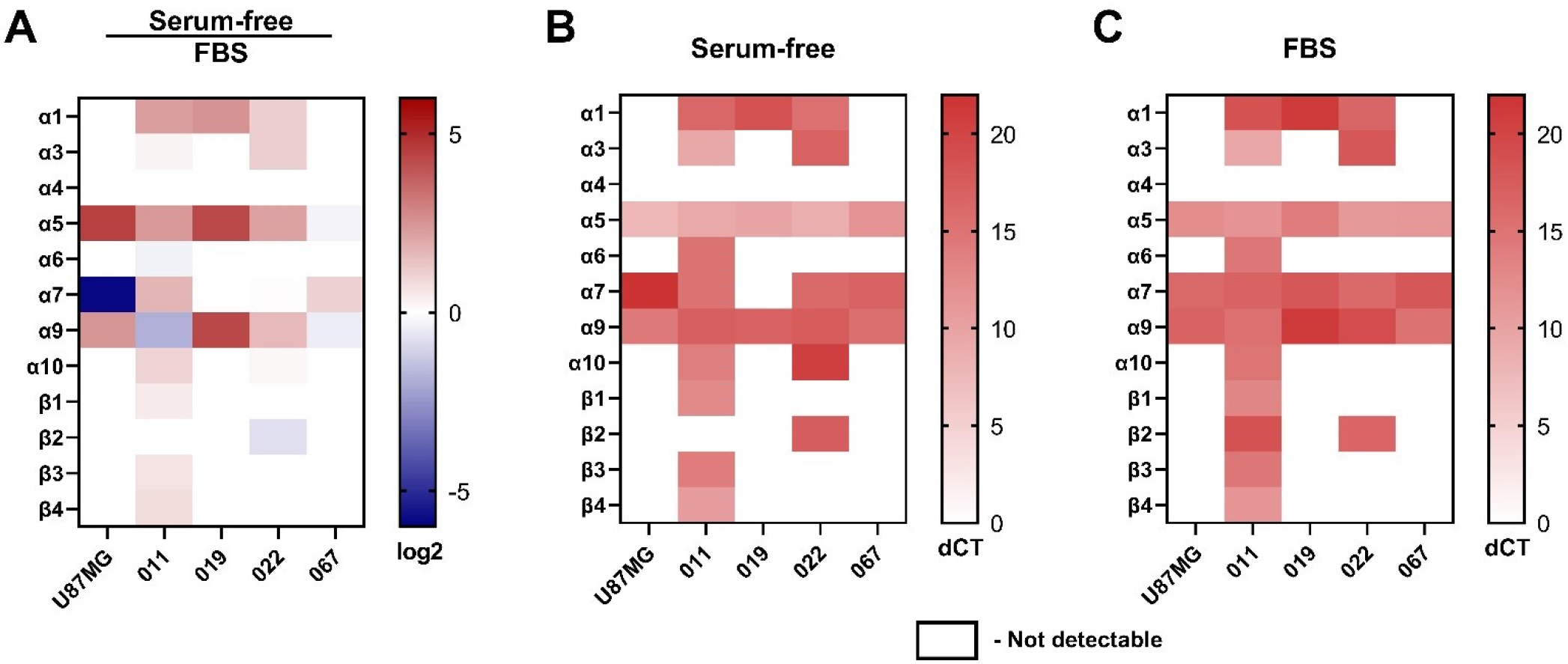
Relative qPCR analysis of nAChR subunits genes expressed in primary GBM lines and model line U87MG. **A**: Relative gene expression was calculated as base 2 logarithm intensity values (Serum-free samples over FBS samples), and represented as a heatmap; **B**: dCT values of nAChR subunits genes were normalized over the 18S level in the corresponding cell lines cultured in serum-free conditions; **C**: dCT values of nAChR subunits genes were normalized over the 18S level in the corresponding cell lines cultured in FBS-supplemented medium.

### Surface α7 nAChR expression confirmed by Alexa 555 labeled α-Bgt

To visualize nAChRs on cell membrane a labeling with 50 nM Alexa Fluor 555-α-bungarotoxin (α-Bgt-AF555) was used. Multi-banded krait venom α-neurotoxin α-Bgt is a competitive antagonist of α7, α9 and muscle-type nAChRs which binds to these receptors with a high affinity. Intensity of α-Bgt-AF555 staining significantly exceeded background level (Fig. 4 A). To confirm the specificity of the α-Bgt labeling protocol 20-fold molar excess of d-tubocurarine (d-TC), another competitive antagonist of nAChRs, was added and significant reduction of α-Bgt labeling was observed (Fig. 4 B).

**Figure 4.**
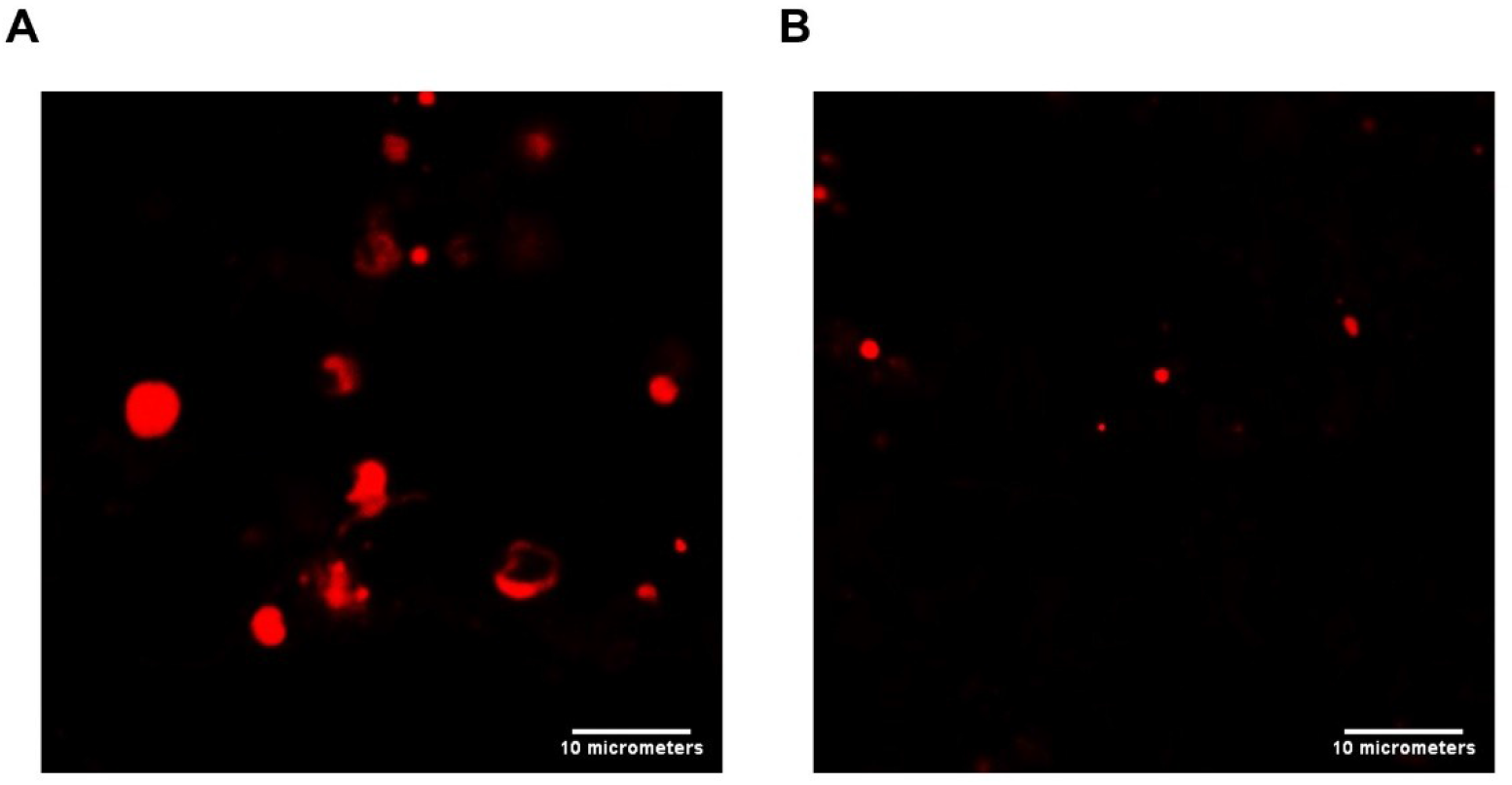
Confocal laser scanning microscopy. **A**: Mesenchymal GBM cells 022 incubated with the α-Bgt-Alexa 555 fluorescent label; **B**: GBM cells co-incubated with α-Bgt-Alexa 555 and excess amount of d-TC without fluorescent label.

### Calcium imaging and patch-clamp electrophysiology of nAChR functional activity

Fluorescent single-cell calcium imaging was performed to assess whether nAChRs gene expression observed by RNA-Seq database analysis from Ivy GAP and RT-PCR correlated with expression of functional nAChRs. Since mRNA expression of calcium-conducting α1, α7 and α9 subunits was detected, functional tests were performed with ligands specific to the respective nAChR subtypes (Fig. 5).

**Figure 5.**
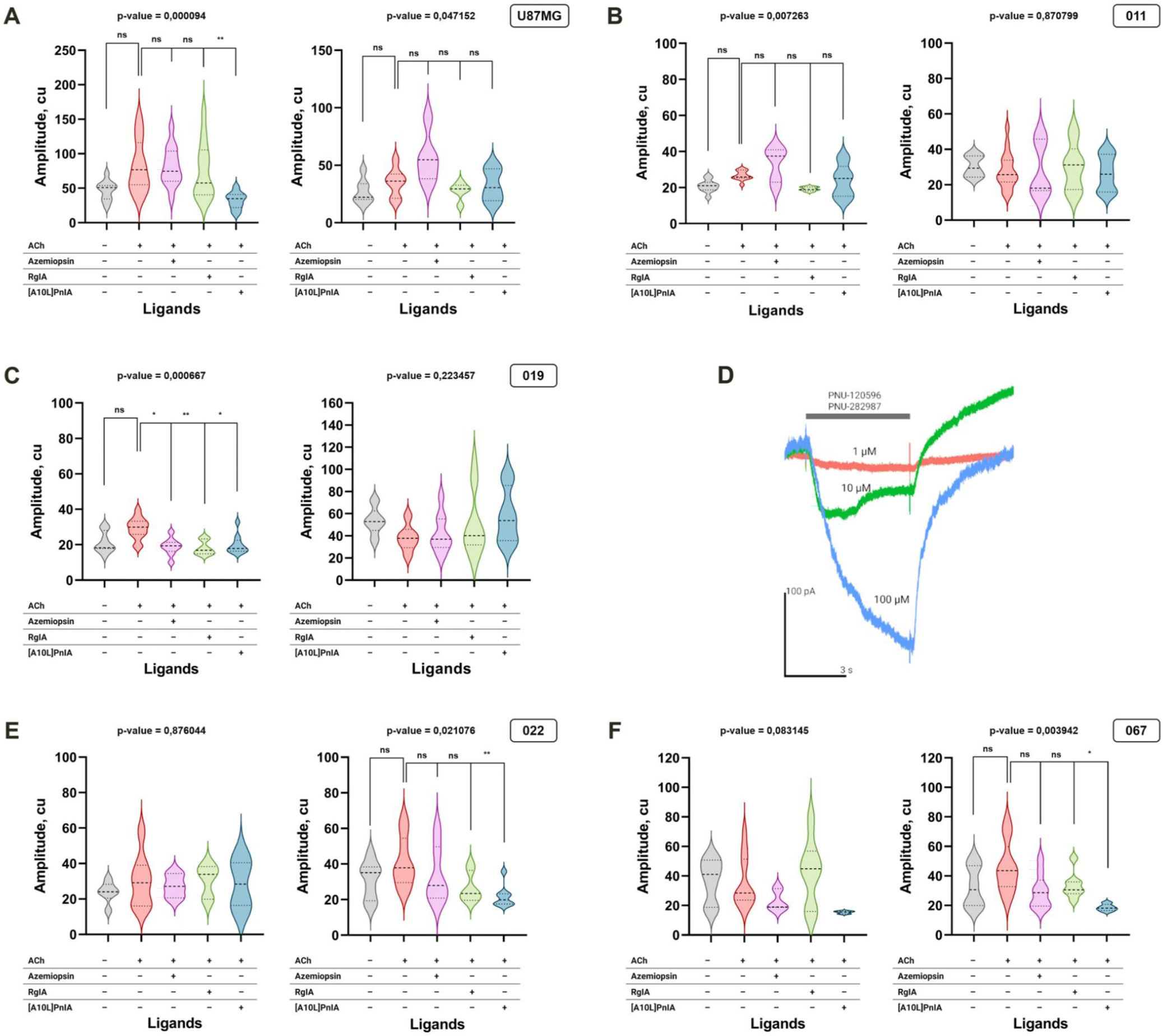
Effects of nicotinic ligands on the intracellular Ca^2+^ concentration in GBM. Cells were loaded with Fluo-4 fluorescent calcium indicator. Left diagram of each panel shows acetylcholine-evoked Ca^2+^ rise of cells grown on FBS-supplemented medium, right diagram of the panel demonstrates responses of serum-free (NS-21/EGF/bFGF-supplemented) medium. **A**: U87MG glioblastoma model line responded to acetylcholine application when cultured on FBS-supplemented medium. This response was significantly reduced by α7 nAChR selective antagonist [A10L]PnIA but not by azemiopsin and RgIA; **B**: Proneuronal line 011 did not demonstrate significant nAChR activation on both culture medium types; **C**: Pro-neuronal GBM cell line 019 demonstrated strong evidence of muscle, α7 and α9 nAChR functioning on FBS-supplemented but not the serum-free medium; **D**: The same 019 line has shown concentration-dependent ion currents in whole-cell patch-clamp experiment (FBS-supplemented medium) in response to mixture of α7 nAChR-selective agonist and positive allosteric modulator; **E**: Mesenchymal GBM primary cell line cultured on serum-free medium demonstrated inhibition of acetylcholine-evoked calcium rise by α7 nAChR-selective inhibitor [A10L]PnIA, whereas the FBS-supplemented cells did not show significant responses; **F**: Another mesenchymal primary GBM cell line have shown responses similar to 022 line that were inhibited by [A10L]PnIA in serum-free culturing conditions.

All cell lines were responsive to the application of 10 μM acetylcholine (ACh). To demonstrate the specificity of nAChR activation, nAChR inhibitors were used in combination with ACh. GBM cells were preincubated with azemiopsin, a selective competitive antagonist of muscle-type nAChR, and with two α-conotoxins, [A10L]PnIA, which selectively blocks α7 nAChR, and RgIA, which inhibits α9 nAChR. A functionally active α7 nAChR was detected on the U87MG model line cultivated on FBS-supplemented medium. As shown in Fig. 5C one of the pro-neuronal cell lines (line 019) grown on a serum-supplemented medium expresses α7, α9 receptors and muscle nAChR. This line grown on serum-supplemented medium adhered well to the coverslip used in patch-clamp technique and whole-cell dosedependent currents were recorded in response to α7 nAChR-selective superagonist PNU-282987 in the presence of positive allosteric modulator PNU-120596, thus confirming cell-surface expression of functionally active α7 nAChR (Fig. 5 D). However, no functionally active receptors were observed on the same line when cultivated on a serum-free medium with growth factors. On the contrary, both primary cultures of the mesenchymal subtype, which were grown on a serum-free medium with growth factors, expressed α7 nAChR, whereas no nAChR were detected when these lines were cultivated on serum-supplemented medium (Fig 5 E and F).

### GBM cell viability after exposure to nAChR agonists and antagonists

The effect of nAChR ligands on GBM cell proliferation was studied using resazurin-based colorimetric method (AlamarBlue). Expression of the calcium-conducting nAChRs containing α1, α7, and α9 subunits was detected at the protein level by fluorescence microscopy and functional tests (calcium imaging and patch-clamp) with specific ligands. Noteworthy, it was reported that GBM tumor stem cells showed strong differences under serum and serum-free cultivation conditions [20]. Cell culturing conditions play an important role in obtaining representative data because the response of cells to different agents can vary. To evaluate the effect of serum on GBM cell proliferation in response to nAChR ligands, the tests were performed on two media variants: serum-free and with FBS addition.

It was found that exposure to varying concentrations of non-selective nAChR agonist epibatidine did not influence cell proliferation (Fig. 6). Addition of α7 nAChR antagonist [A10L]PnIA along with epibatidine led to an increase in cell proliferation both with FBS (Fig. 6 A, E, G, E) and in serum-free medium in 4 of 5 analyzed lines: 011, 022, 067, and U87MG (Fig. 6 F, H, J). Line 067 cultured on FBS-supplemented medium exhibited increased proliferation in response to exposure to the muscle nAChR antagonist azemiopsin in presence of epibatidine (Fig. 6 G). Azemiopsin also increased proliferation of 022 and U87MG cell lines grown on serum-free medium (Fig. 6 F and J).

**Figure 6.**
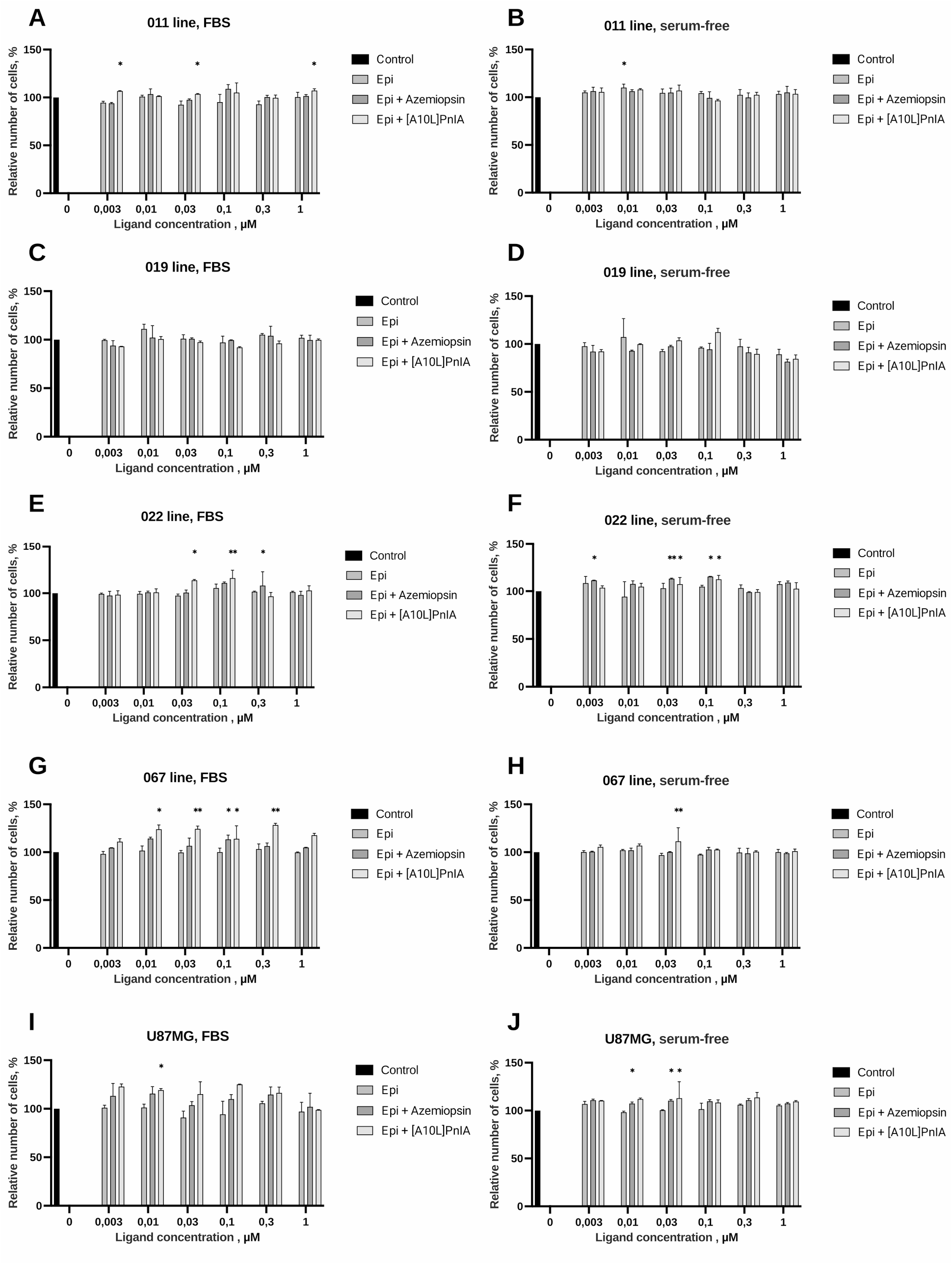
AlamarBlue cell viability assay results of nAChR ligands influence on cell proliferation in FBS and serum-free medium. Values are normalized to control without adding ligands. **A**: Pro-neuronal primary GBM cell line 011 grown on FBS-supplemented medium demonstrated increase in cell proliferation under the influence of [A10L]PnIA in the presence of epibatidine; **B**: epibatidine (10 nM) significantly increased proliferation of pro-neuronal primary GBM cell line 011 grown on serum-free medium; **C**: No significant effects were detected in case of pro-neuronal primary GBM cell line 019 grown on FBS-supplemented medium; **D**: No significant effects were detected in case of pro-neuronal primary GBM cell line 019 grown on serum-free medium; **E**: Mesenchymal primary GBM line 022 grown on FBS-supplemented medium increased proliferation in response to [A10L]PnIA in combination with 30 nM and 100 nM epibatidine. Azemiopsin in combination with 300 nM epibatidine slightly increased the proliferation; **F**: Mesenchymal primary GBM line 022 grown on serum-free medium increased proliferation in response to [A10L]PnIA in combination with 30 nM and 100 nM epibatidine. Azemiopsin in combination with 3 nM epibatidine slightly increased the proliferation; **G**: Mesenchymal primary GBM line 067 grown on FBS-supplemented medium increased proliferation in response to [A10L]PnIA in combination with 10 nM – 300 nM epibatidine. Azemiopsin in combination with 300 nM epibatidine slightly increased the proliferation; **H**: [A10L]PnIA in combination with 30 nM epibatidine increased the proliferation of mesenchymal primary GBM line 067 grown on serum-free medium; **I**: Model GBM line U87MG grown on FBS-supplemented medium increased proliferation in response to combination of [A10L]PnIA and 10 nM epibatidine; **J**: U87MG grown on serum-free medium increased proliferation in response to azemiopsin in combination with epibatidine at 10 and 30 nM.

It should be noted that pro-neuronal GBM from patients 011 and 019 acted differently in response to the nAChR antagonists application. Patient 011 cells increased proliferation (especially on medium containing serum), whereas in 019 cells the cell proliferation level did not change in all media options (Figure 6 A-D). Cells of the mesenchymal subtype (022 and 067), which are characterized by a high level of stemness and a more aggressive phenotype, also showed different results. The observed difference can be explained by the extreme heterogeneity of GBM cells, meaning that the cells belonging to the same phenotype may have different properties and respond differently to the action of different agents.

Thus, almost no direct influence of epibatidine on GBM cells proliferation was detected but addition of nAChR inhibitors surprisingly showed pro-proliferative effects. The second AlamarBlue test was deployed to study cells incubated with the addition of α1, α7, and α9 nAChR antagonists but without the agonist in two media: with and without FBS (Fig. 7).

**Figure 7.**
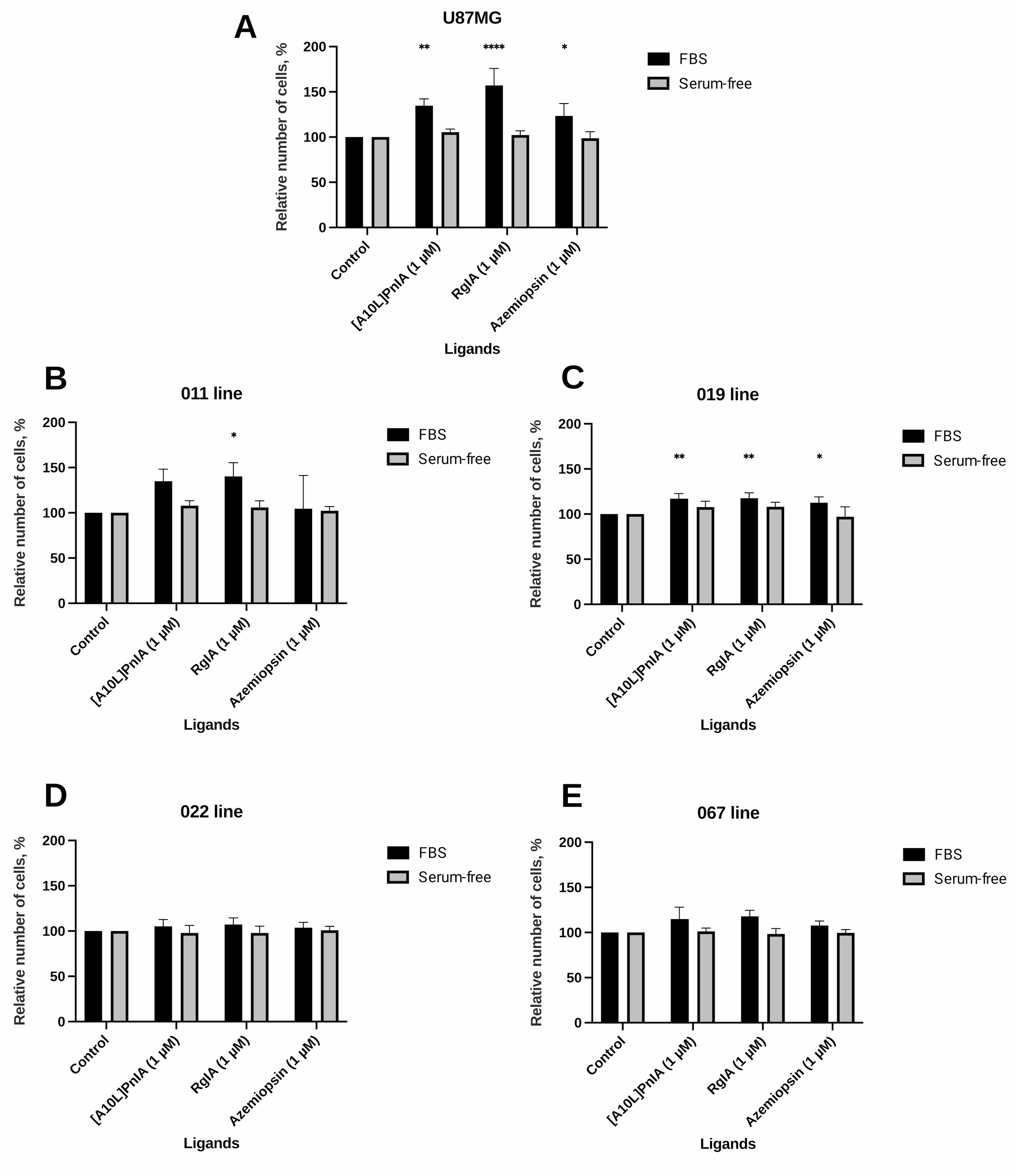
AlamarBlue cell viability assay study of the effects of nicotinic antagonists exposure on *in vitro* GBM cell proliferation under different cultivation conditions. Values are normalized to control without adding ligands. **A**: Model GBM line U87MG increased proliferation in response to [A10L]PnIA, RgIA and azemiopsin while being grown on FBS-supplemented but not on the serum-free medium; **B**: Proneuronal GBM cell line 011 significantly increased proliferation in response to α9 nAChR inhibitor RgIA in FBS-supplemented but not the serum-free medium; **C**: Pro-neuronal GBM cell line 019 increased proliferation in response to [A10L]PnIA, RgIA and azemiopsin while being grown on FBS-supplemented but not on the serum-free medium; **D**: No significant effects of nAChR antagonists were detected for the mesenchymal GBM cell line 022 neither on FBS-supplemented nor on serum free-medium; **E**: No significant effects of nAChR antagonists were detected for the mesenchymal GBM cell line 067 neither on FBS-supplemented nor on serum free-medium.

The U87MG cell line and the pro-neuronal lines (011 and 019) increased proliferation in response to exposure to nAChR antagonists in the FBS-supplemented medium, whereas under serum-free conditions cell proliferation remained at the control level (Fig. 7 A-C). U87MG cell line showed an increased cell proliferation in response to exposure to all nAChR antagonists used, moreover, the greatest increase in proliferation occurred in response to the α9 nAChR antagonist α-conotoxin RgIA (Fig. 7 A). Line 011 responded significantly only to this antagonist, but line 019 increased proliferation when exposed to azemiopsin and α-conotoxin [A10L]PnIA, antagonists of muscle and α7 nAChRs (Fig. 7 B-C). Cells of the mesenchymal subtype lines (022 and 067) showed no significant changes in proliferation in both media variants (Fig. 7 D-E).

Fetal bovine serum (FBS) used to supplement cell culture medium is basically the blood serum and contains up to 10–100 μM of choline [26]. Two explanations for the results have been proposed: i) the effect of choline contained in serum increases cell proliferation; ii) cell cultivation in medium with FBS leads to an increase in cell differentiation due to a lower concentration of growth factors then in serum-free medium, supplemented with EGF and bFGF.

To get understanding of choline influence on nAChR antagonists-mediated increase in proliferation, two more AlamarBlue tests were performed. In the first one, serum-free medium with the addition of growth factors (bFGF and EGF) and a neuronal vitality supplement (NS-21) was used. Varying concentrations of nAChR agonist (in this case choline) in combination with antagonists as in the previous experiment (Fig. 6). The model line U87MG showed no significant changes in cell proliferation (Fig. 8 A). Line 011 showed no significant changes in cell proliferation (Fig. 8 B), whereas line 019 showed increased proliferation in response to all ligands at 1 μM choline concentration, as well as an increased proliferation in response to azemiopsin, an α1 nAChR antagonist at 10 μM choline concentration (Fig. 8 C). Line 022 showed no significant changes in cell proliferation (Fig. 8 D), whereas line 067 showed an increase in cell proliferation at choline concentrations from 0.1 to 10 μM in response to α7 and α9 nAChR antagonists (Fig. 8 E). It is worth noting that of all the lines studied, line 067 is the most aggressive and resistant to therapy. These results may confirm that the action of choline present in FBS, can influence cell growth through persistent activation of muscle, α7 and α9 nAChRs.

**Figure 8.**
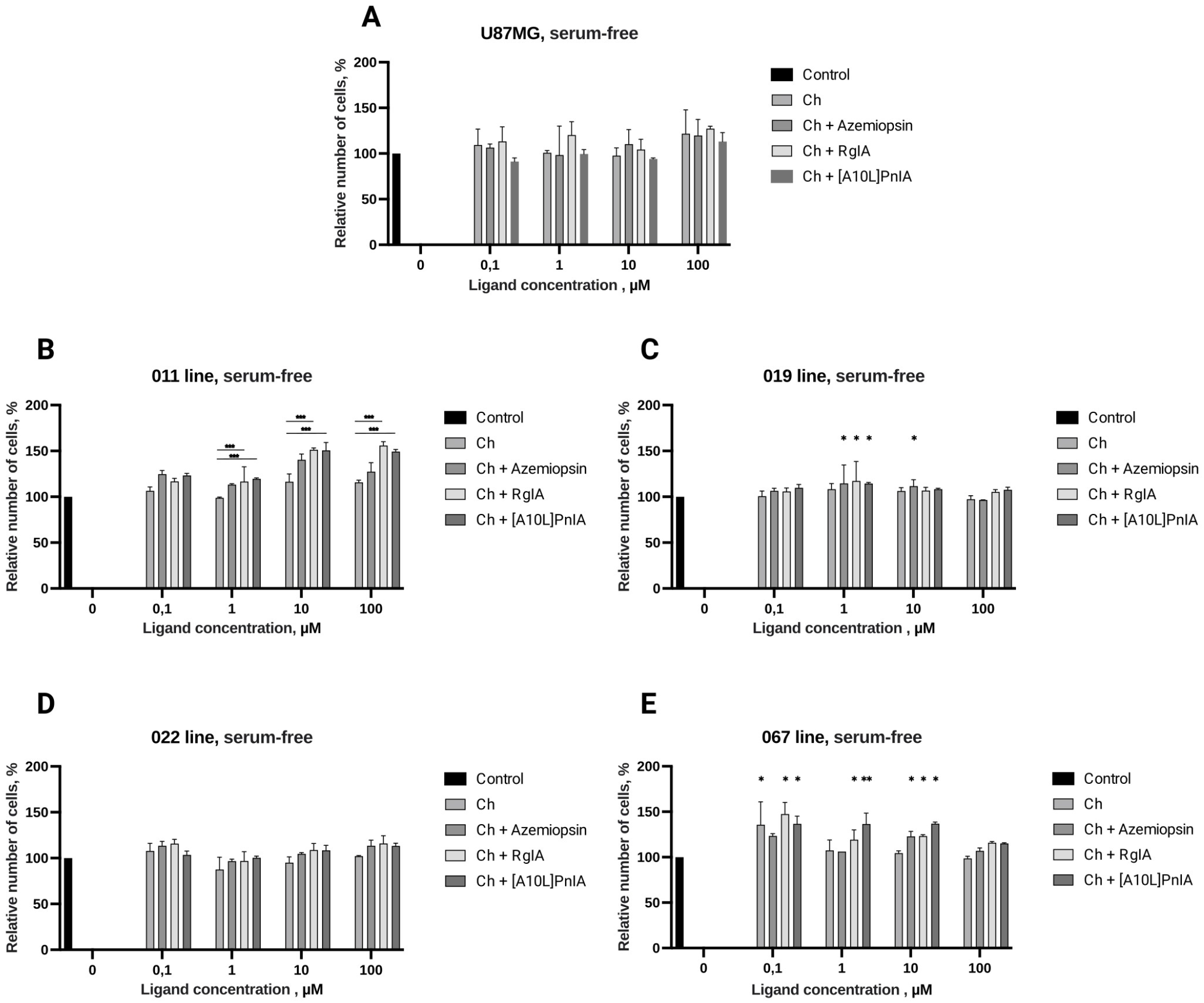
AlamarBlue cell viability study of the effects of nicotinic agonist choline and in the presence of various antagonists on GBM cells in serum-free medium. Values are normalized to control without adding the ligands. **A**: No significant effects of choline and its combination with nAChR antagonists was found for U87MG GBM model line kept in serum-free medium; **B**: Addition of [A10L]PnIA and RgIA in combination with 1-100 μM of choline significantly increased proliferation of pro-neuronal GBM line 011; **C**: Significant increase in the proliferation of pro-neuronal GBM line is observed under the influence of mixture nAChR inhibitor (azemiopsin, [A10L]PnIA or RgIA) with 1 μM choline; **D**: No significant effects of choline and its combination with nAChR antagonists was found for mesenchymal GBM line 022; **E**: Choline (100 nM) stimulated proliferation of mesenchymal GBM line 067. This stimulatory effect was diminished by azemiopsin. Choline at concentrations 1–100 μM did not stimulate cell proliferation but all three selective nAChR inhibitors increased proliferation in combination with 10 μM choline.

Line 067 is the most aggressive among the tested ones and it increased proliferation in response to the action of muscle, α7 and α9 nAChRs antagonists over a wide range of choline concentrations. Growth factors influence on the nAChR-solicited effect on proliferation was further investigated using the 067 line under a various conditions. Tests with and without an agonist (epibatidine) were performed (Fig. 9). Line 067 showed no significant changes in cell proliferation in the NS-21-supplemented medium without the addition of bFGF and EGF (Fig. 9 A). There was a slight increase in cell proliferation in the medium without EGF but with the addition of bFGF and NS-21 in response to the α7 nAChR antagonist (Fig. 9 A-B). Moreover, there was a strong decrease in cell proliferation in response to epibatidine in empty medium containing no growth factors. Muscle nAChR antagonist azemiopsin diminished the response to low concentrations of epibatidine (0.03 and 0.1 mM Fig. 9 C). Interestingly, α7 nAChR antagonist [A10L]PnIA rescued GBM cells from growth suppression by 0.03, 0.1, 0.3, and 1 mM epibatidine. These results may indicate that in the absence of growth factors, the activation of α7 and α1* (muscle) nAChR significantly reduces cell proliferation. Cell proliferation of line 067 was significantly increased in response to the α7 and α9 nAChR antagonists (no external agonists being applied) in medium without EGF but with the addition of bFGF and NS-21 (Fig. 9 E). However, there was no change in cell proliferation in empty medium (Fig. 9 F). A slight increase in proliferation in response to the α1 nAChR antagonist azemiopsin was observed in NS-21-supplemented medium without EGF and bFGF (Fig. 9 D).

**Figure 9.**
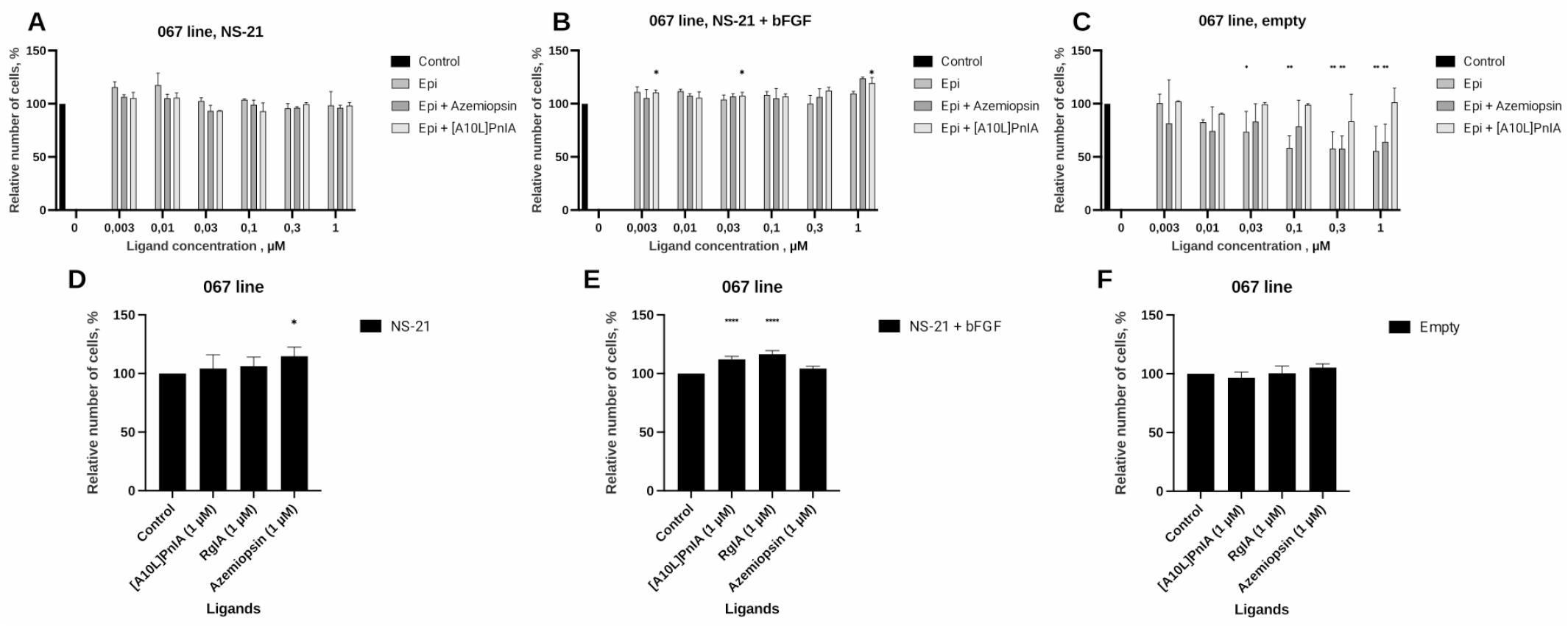
AlamarBlue cell viability assay of the effects of nicotinic ligands *in vitro* exposure of the 067 line GBM cell under various cultivation conditions. Values are normalized to control without adding ligands. **A**: No significant effects of epibatidine and of its combination with nAChR antagonists was found on cells kept in NS-21-supplemented serum-free medium without addition of EGF and bFGF; **B**: Addition of bFGF to the NS-21-supplemented medium leads to emergence of pro-proliferative effect of a combination of [A10L]PnIA with 3 nM, 30 nM and 1 μM of epibatidine; **C**: Proliferation of GBM cell line 067 kept in plain DMEM/F12 medium was significantly reduced by epibatidine in concentrationdependent manner. Azemiopsin and [A10L]PnIA reduced the effect of epibatidine, suggesting the involvement of muscle and α7 nAChRs, respectively; **D**: On NS-21-supplemented medium azemiopsin significantly stimulated proliferation of 067 cells even in the absence of externally added agonists; **E**: Azemiopsin had no effect whereas [A10L]PnIA and RgIA significantly stimulated proliferation of the cells on NS-21/bFGF-supplemented medium; **F**: No effect of nAChR antagonists were detected on plain DMEM/F12 medium.

## DISCUSSION

In recent years, a wide range of methods have been proposed in the study of GBM, which made it possible to understand the mechanisms of tumor development. Cancer cell lines have been the standard both for studying the biology of human tumors and as preclinical screening models of potential therapeutic agents. However, it is becoming increasingly clear that phenotypic characteristics and a variety of genetic aberrations found in cancer cell lines and repeatedly passaged *in vitro* often bear little resemblance to those found in the corresponding human primary tumor. This may have led to some important misinterpretations regarding the significance of aberrant signaling pathways within cell lines as compared to primary tumors. GBM tumor cell populations have a variety of functional and molecular biological features, which makes the tumor extremely heterogeneous [18]. In particular, glioblastoma stem cells (GSCs), which are characterized by increased resistance to chemotherapy and radiotherapy, are present in the overall cell population [28].

In current work, the effect of the cultivation methods on the nAChR expression was investigated. The presence of nAChR subunit gene expression in primary cultures of samples 011, 019, 067, and 022 at the mRNA level was demonstrated. Under different cell culture conditions (FBS-supplemented vs. NS-21/EGF/bFGF) the primary neurosphere GBM cultures from different human sources have shown distinct profile of the nAChR genes expression (Fig. 3). Noteworthy, α5 and α9 nAChR subunits were detected in all GBM lines tested. GBM line 011 expressed β2 nAChR subunits gene exclusively when cultured on FBS-supplemented medium, while the α9 nAChR subunit was significantly increased in the FBS-supplemented in contrast to serum-free medium. Lines 011, 019 and 022 showed a significant increase in the muscle nAChR α1 subunit gene. Gene encoding α7 nAChR subunit was reduced in serum-free as compared to the FBS-supplemented medium in U87MG model cell line but in 019 line the expression of the same nAChR gene was observed only on FBS-supplemented medium and other lines expressed this receptor independent on culturing conditions. Overall, mesenchymal lines 022 and 067 demonstrated fewer differences in nAChR genes expression dependent on FBS or serum-free medium than pro-neuronal lines 011 and 019. Mesenchymal line 022 synthesized the functionally active nAChR which was confirmed by specific fluorescent ligand binding, namely Alexa Fluor 555 α-bungarotoxin (Fig. 4), showing that nAChR genes expression leads to the respective protein production by GBM cells.

Functioning of nAChRs in GBM cell cultures were studied using selective antagonists –α-conotoxins [A10L]PnIA and RgIA which bind preferably to α7 and α9 nAChRs, respectively. Neurotoxin azemiopsin from *Azemiops feae* viper venom was used to selectively detect muscle (α1β1δε) nAChR. The activity of the receptors was monitored using fluorescent Ca^2+^ imaging. Acetylcholine application to cells evokes a rise of intracellular Ca^2+^ that is detected as fluorescence intensity increase (Fig. 4). If this rise in calcium is depressed by selective antagonist it can be interpreted as mediated (at least in part) by the respective nAChR subtype. Interestingly, U87MG cell line showed a significant depression of the acetylcholine-evoked calcium rise by α7 nAChR-targeted [A10L]PnIA and only on cells cultured on FBS-supplemented medium (Fig. 4 A), which is in a good agreement with RT-PCR data showing decline in α7 nAChR gene expression in serum-free conditions. However, no significant evidence of functioning muscle and α9 nAChR was found in cells grown on FBS or serum-free medium.

One of the pro-neuronal GBM cell lines, namely 011 did not show signs of acetylcholine-evoked calcium rise that can be inhibited by muscle, α7 or α9 nAChR inhibitors, suggesting limited presence of functioning receptors of these subtypes. It should be noted that for the cells grown on the FBS-supplemented medium Kruskal-Wallis test detected significant (p < 0.01) differences in calcium rise amplitudes for control and inhibitor-treated cells (Fig. 4 B, left diagram). Pairwise Dunn’s test, however, did not detect significant differences between groups, suggesting that the effect of selective inhibitors, if present, is of a limited size. The other pro-neuronal GBM line tested, 019, demonstrated strong evidence of muscle, α7 and α9 nAChR when cultured on FBS-supplemented medium (Fig. 4 C). The presence of α7 nAChR was further supported by patch-clamp recordings of whole-cell currents elicited by selective α7 nAChR agonist in combination with selective α7 nAChR positive allosteric modulator (Fig. 4 D). Despite being detected by RT-PCR in 019 cells grown on both types of medium, no significant effects of selective inhibitors were detected in calcium imaging experiments on cells cultured in serum-free conditions (Fig. 4 C, right panel), suggesting significant influence of culture conditions on the receptors translation or membrane transport. In contrast to 011 and 019 pro-neuronal GBM lines, both mesenchymal lines (022 and 067) have shown strong evidence of α7 nAChR functioning only when cultured on serum-free medium (Fig. 4 E and F, respectively). These results demonstrate the variability of the GBM cell lines in respect to functional nAChR surface expression.

The immortalized U87MG cell line has been used in many laboratories worldwide, but due to genetic drift in the FBS-supplemented medium and successive passages, the cells with the highest proliferative potential are selected, reducing the genetic heterogeneity inherent to the original tumors, which can affect the reproducibility of studies [29]. Primary cell cultures obtained from patients are more representative models, but the use of media containing fetal bovine serum (FBS) can lead to loss of the GSCs subpopulation [29], making the tumor homogeneous. Among other things, the presence of GSCs is closely linked to the formation of spheroids in tumor cell cultures, called neurospheres of neurally originated tumors [30]. It was previously shown that glioma cells cultured in the medium with serum or in the serum-free medium showed profound biological differences *in vitro* [20]. One approach to prevent loss of tumor heterogeneity and culturing the cells as neurospheres is the use of a serum-free liquid medium containing basic fibroblast growth factor (bFGF), epidermal growth factor (EGF) and a neuronal viability supplement (B27 or NS-21) [31]. In our paper the interconnection between culturing condition and nAChR gene expression and functioning is described for the first time using nAChR subtype-selective neurotoxins as a research instrument.

It was recently reported by Pucci et al. that nicotine and choline increased the proliferation of glioblastoma model lines U87MG and GBM5 [17]. In this report U87MG line was cultured in the FBS-supplemented medium and GBM5 was kept in serum-free conditions supplemented with Neurobasal and B27 additives. Interestingly, Neurobasal supplement contains choline at concentration of 28.6 μM. Choline in blood serum is estimated to be in range of 7.1 – 28.9 μM [27]. And some reports estimated choline concentration in blood plasma using LC-MS/MS at the level of 11.4 ng/μL, which corresponds to ~ 100 μM [26], suggesting that in 10% FBS-supplemented medium and in 1:1 DMEM/F12 with Neurobasal mixture choline levels can reach the concentration of several μM.

Choline is metabolic precursor of the major endogenous cholinergic neurotransmitter acetylcholine and plays an important role in the nAChR functioning. It has been shown to act as agonist of the muscle, α7 and α9 nAChR and a modulator of the α3β4 nAChR [23,32,33]. In current article the effects of nicotinic agonists on various GBM cell lines were investigated with respect to cell culturing conditions. To exclude the possibility of choline influencing the results, serum-free medium was based on Neuro Brew^™^ NS-21 supplement (reformulated variant of B27) that does not contain choline.

In current study, no significant effect of nAChR agonist epibatidine was detected over a wide range of concentrations (3 nM – 1 μM) on the U87MG and primary GBM neurosphere cultures (Fig. 6). Surprisingly, the application of nAChR antagonists along with epibatidine in some cases stimulated proliferation. Modified α-conotoxin [A10L]PnIA which selectively inhibits α7 nAChR stimulated proliferation of pro-neuronal 011 line grown on the FBS-supplemented medium (Fig. 6 A) while no effects of antagonists were detected in the serum-free medium (Fig. 6 B). No effect of nAChR ligands on proliferation was detected with the pro-neuronal 019 line (Fig. 6 C and D) despite this line clearly shows presence of functional nAChRs in calcium imaging and patch-clamp experiments. The proliferation of mesenchymal lines 022 and 067 and model line U87MG cultured on FBS-supplemented and serum-free media were stimulated by [A10L]PnIA suggesting the possible role of α7 nAChR which is in a good agreement with calcium imaging and RT-PCR results (Fig. 6 E-J). The selective muscle nAChR inhibitor azemiopsin stimulated 022 mesenchymal line and U87MG model line in both types of medium (FBS and serum-free, see Fig. 6 E, F, I and J) but for mesenchymal line 067 its effect was detected solely in FBS-supplemented medium (Fig. 6 H).

It is not clear why epibatidine being a potent nAChR agonist did not stimulate GBM cells proliferation (apart from subtle effect on pro-neuronal 011 line in serum-free medium, see Fig. 6 B). Even more surprisingly, exactly opposite, namely the inhibition of nAChRs, stimulated the proliferation. The antagonists influence on proliferation was also studied in the absence of externally added agonists (Fig. 7). Strikingly, on FBS-supplemented medium U87MG model line and two pro-neuronal GBM primary neurosphere cultures 011 and 019 have demonstrated significant increase in proliferation in response to α9 nAChR-selective antagonist RgIA (Fig. 7 A-C). U87MG and 019 lines also increased proliferation in response to muscle and α7 nAChR-selective antagonists in FBS-supplemented medium (Fig. 7 A and C). No significant nAChR antagonist-stimulated proliferation was detected on serum-free medium for these lines. Mesenchymal GBM spheroid primary cultures 022 and 067 did not increased proliferation in response to nAChR antagonists on either FBS-supplemented or serum-free medium (Fig. 7 D and E).

Since nAChR inhibitors-provoked proliferation tend to emerge (although not exclusively) in FBS-supplemented medium which by design contains choline as a component of blood serum, possible role of choline was studied by external addition of its varying concentrations to the serum-free medium (Fig. 8). No strong evidence of choline provoking cell proliferation was detected on the serum-free medium for U87MG, 011, 019 and 022 lines (Fig. 8 A-D). There was some indication of 100 nM choline-stimulated proliferation of 067 mesenchymal line (Fig. 8 E). However, similarly to results of epibatidine experiment (Fig. 6), the addition of selective antagonists along with choline provoked proliferation of 011, 019 and 067 lines (Fig. 8 B, C and E). Note, that no antagonist-stimulated proliferation has been detected on the same serum-free medium when no choline was added (Fig. 7) suggesting that some basal nAChR activation by choline is needed for nAChR antagonist-stimulated proliferation.

Serum-free medium used in the current study contains EGF and bFGF. EGF has previously been shown to support growth and tumor spreading [34]. In particular, EGF is secreted by tumor-associated macrophages and microglia. EGF secretion plays a role in the mechanisms of healing and regeneration of brain tissues, determining the migration of microglia to GBM lesions [35]. EGF influences glial-mesenchymal transition and promotes microevolution of GBM malignancy, enhances the invasive potential of GBM cells and their ability to penetrate healthy tissues [36]. Thus, the conditions for growing cell cultures determine the heterogeneity of the cell population and their microevolution even in the absence of an active microenvironment. To account for the possible influence of growth factor combinations on the nAChR antagonists-stimulated proliferation, the 067 line was grown on NS-21-supplemented serum-free medium, bFGF-supplemented serum-free medium (NS-21/bFGF) or plain serum-free DMEM/F12 (Fig. 9). Notably, on medium supplemented solely with NS-21 no significant effects of epibatidine or α7 and α9 nAChR antagonists in presence of epibatidine have been detected (Fig. 9 A). Cells cultured in presence of bFGF were stimulated by α7 nAChR antagonist [A10L]PnIA (in mixture with epibatidine) but no effects of epibatidine itself on the proliferation have been detected (Fig. 9 B). Interestingly, on plain serum-free DMEM/F12 epibatidine significantly reduced cells proliferation and its effect was suppresed by muscle nAChR-selective inhibitor azemiopsin and by [A10L]PnIA (Fig. 9 C).

Somewhat unexpectedly azemiopsin significantly stimulated proliferation of 067 cells grown on NS-21-supplemented serum-free medium whithout EGF and bFGF (Fig. 9 D). Addition of bFGF to the medium diminished azemiopsin-stimulated proliferation and promoted [A10L]PnIA and RgIA effects, which might indicate that bFGF enhance α7 and α9 nAChR expression or function but inhibits expression of muscle nAChR (Fig. 9 E). No pro-proliferative effect of antagonists was detected on plain DMEM/F12 serum-free medium (Fig. 9 F), suggesting that despite functional expression of muscle and α7 nAChR (compare Fig. 9 C and F), the antagonists-stimulated effects are dependent on the receptor activation by some agonist. NS-21 on which Neuro Brew^™^ used in the current study is based, contains ethanolamine [37]. To our knowledge, ethanolamine does not activate human nAChRs but is a predecessor of the choline and acetylcholine in the biosynthetic pathways which might explain significant stimulation of cell growth by α7 and α9 nAChR antagonists on serum-free medium containing Neuro Brew^™^(Fig. 9).

## CONCLUSIONS

In this paper we describe highly variable consequences of nicotinic acetylcholine receptors (nAChR) activation and inhibition on primary glioblastoma multiforme (GBM) spheroid cultures and a model line U87MG. Not only nAChR gene expression varies between different patients but actual receptor functioning and cell proliferation depends on nAChR agonists and antagonists. The results reported here may be helpful in subsequent studies design focusing on cell culturing conditions and interpretation of nAChR ligands effects. It is also possible to to monitor blood plasma and cerebrospinal fluid choline level for better interpretation of clinical studies focused on the role of tobacco smoking in glioma patients.

## DATA ACCESSIBILITY

All data reported in the manuscript are available from the corresponding author on demand.

## AUTHOR CONTRIBUTIONS

D.K. conceived the study, obtained the funding, conducted the patch clamp experiment, supervised the calcium imaging and wrote the paper; D.M. performed the AlamarBlue assays; E.G. performed the calcium imaging experiments and wrote the paper; I.K. performed the experiments; M.M analyzed the transcriptomic data and wrote the paper; N.A. supervised the cell culturing and AlamarBlue experiments; V.T. edited the paper; Y.R. performed confocal fluorescence microscopy.

## ACKNOWLEDGEMENTS

Authors are grateful to Dr. Marat Pavlyukov for valuable comments.

