## Supplemental Table 1 for "Patient-derived glioblastoma neurosphere cultures differentially express nicotinic acetylcholine receptors depending on ambient choline"

Table 1. RT-PCR oligonucleotide primers used for detection of *Homo sapiens* nAChR genes expression.

| Gene code | Sequence |
|---|---|
| CHRNA1 | forward<br>5’ – GGCTCCGAACATGAGACCCG – 3’ |
| | reverse<br>5’ – CCACTCCTCAGACGCATTG – 3’ |
| CHRNA3 | forward<br>5’ – GTGGATGAAGTAAACCAGATCATGGAG – 3’ |
| | reverse<br>5’ – GGAACCGAACTTCATGGTACAGTTTTG – 3’ |
| CHRNA4 | forward<br>5’ – CCTCGGCCTGTCCATCGCTCA – 3’ |
| | reverse<br>5’ – AAGACGGTGAGCGACAGCAGC – 3’ |
| CHRNA5 | forward<br>5’ – ACTGTCACCTGGACTCCACCG – 3’ |
| | reverse<br>5’ – AACAGCTGTCGGTTCTGTTTCCTTTG – 3’ |
| CHRNA6 | forward<br>5’ – TGTGGGCTGTGCAACTGAGGAG – 3’ |
| | reverse<br>5’ – CAATGTCGGGCTTCCAAATCTTATC– 3’ |
| CHRNA7 | forward<br>5’ – CCCGGCAAGAGGAGTGAAAGGT – 3’ |
| | reverse<br>5’ – ACCGAGAGGCCCACGATGATC – 3’ |
| CHRNA9 | forward<br>5’ – AGAGCCTGTGAACACCAATGTGG – 3’ |
| | reverse<br>5’ – CTCAGAGCAGCAGCCATAGGAG – 3’ |
| CHRNA10 | forward<br>5’ – GATGTAGCAGCCTTCCCGTTCG – 3’ |
| | reverse<br>5’ – AGCAGCAGCGTGAAGGTGACG – 3’ |
| CHRNB1 | forward<br>5’ – GTGTCAGGGTCAGGGTTGGT – 3’ |
| | reverse<br>5’ – CGACGCTAATGTCCAGAGCC – 3’ |
| CHRNB2 | forward<br>5’ – TGTACGAGGTGTCCTTCTATTCCAATG– 3’ |
| | reverse<br>5’ – TGTAGAAGAGCGGCTTGCGGC – 3’ |
| CHRNB3 | forward<br>5’ – CGCCGAAAATGAAGATGCCCTCC – 3’ |
| | reverse |

| | |
|---|---|
| | 5’ – GTCAGCATTTTCAAAGAACTATGTCAGG– 3’ |
| CHRNB4 | forward<br>5’ – CAACAACCTGATCCGCCCAGC – 3’ |
| | reverse<br>5’ – GTTGTAAAGCACGATGTCAGGCAAC – 3’ |
| 18S | forward<br>5’ - GGCCCTGTAATTGGAATGAGTC - 3’ |
| | reverse<br>5’ - CCAAGATCCAACTACGAGCTT - 3’ |
